# 10Kin1day: A bottom-up neuroimaging initiative

**DOI:** 10.1101/509554

**Authors:** Martijn P van den Heuvel, Lianne H Scholtens, Hannelore K van der Burgh, Fredica Agosta, Clara Alloza, Tiffany M C Avancini, Simon Baron-Cohen, Silvia Basaia, Manon JNL Benders, Frauke Beyer, Linda Booij, Kees PJ Braun, Wiepke Cahn, Dara M Cannon, Sandra SM Chan, Eric YH Chen, Cambridge Child Development Project Consortium includes: Auyeung Bonnie; Holt, Rosemary J, Michael V Lombardo, Benedicto Crespo-Facorro, Eveline A Crone, Udo Dannlowski, Sonja MC de Zwarte, Ana M Diaz-Zuluaga, Bruno Dietsche, Stefan Du Plessis, Sarah Durston, Robin Emsley, Geraldo B Filho, Massimo Filippi, Thomas Frodl, Dariusz Gąsecki, Joanna Goc, Martin Gorges, Beata Graff, Dominik Grotegerd, Julie M Hall, Laurena Holleran, Helene J Hopman, Lutz Jäncke, Andreas Jansen, Krzysztof Jodzio, Vasiliy G Kaleda, Jan Kassubek, Shahrzad Kharabian Masouleh, Tilo Kircher, Martijn GJC Koevoets, Vladimir S Kostic, Axel Krug, Stephen M Lawrie, Irina S Lebedeva, Edwin HM Lee, Tristram A Lett, Simon JG Lewis, Franziskus Liem, Carlos Lopez-Jaramillo, Daniel S Margulies, Sebastian Markett, Paulo Marques, Ignacio Martínez-Zalacaín, Colm McDonald, Andrew McIntosh, Genevieve McPhilemy, Susanne L Meinert, José M Menchón, Susan Mérillat, Christian Montag, Pedro S Moreira, Pedro Morgado, David Omar Mothersill, Hans-Peter Müller, Leila Nabulsi, Pablo Najt, Krzysztof Narkiewicz, Patrycja Naumczyk, Sebastiaan WF Neggers, Bob Oranje, Victor Ortiz-Garcia de la Foz, Jiska S Peper, Julian A Pineda Z., Paul E Rasser, Ronny Redlich, Jonathan Repple, Martin Reuter, Pedro GP Rosa, Amber NV Ruigrok, Agnieszka Sabisz, Ulrich Schall, Soraya Seedat, Mauricio H Serpa, Devon Shook, Stavros Skouras, Carles Soriano-Mas, Nuno Sousa, Edyta Szurowska, Alexander S Tomyshev, Diana Tordesillas-Gutierrez, Leonardo Tozzi, Sofie L Valk, Leonard H van den Berg, Theo GM van Erp, Neeltje EM van Haren, Judith JMC van Leeuwen, Arno Villringer, Christiaan Vinkers, Christian Vollmar, Lea Waller, Henrik Walter, Heather C Whalley, Marta Witkowska, Veronica A Witte, Marcus V Zanetti, Rui Zhang, Gary Donohoe, Veronica O’Keane, Siemon C de Lange

## Abstract

We organized 10Kin1day, a pop-up scientific event with the goal to bring together neuroimaging groups from around the world to jointly analyze 10,000+ existing MRI connectivity datasets during a 3-day workshop. In this report, we describe the motivation and principles of 10Kin1day, together with a public release of 8,000+ MRI connectome maps of the human brain.

## Main Text

Ongoing grand-scale projects like the European Human Brain Project (Amunts et al., 2016), the US Brain Initiative (Insel et al., 2013), the Human Connectome Project (Van Essen et al., 2013), the Chinese Brainnetome (Jiang, 2013) and exciting world-wide neuroimaging collaborations such as ENIGMA (Thompson et al., 2017) herald the new era of *big neuroscience*. In conjunction with these major undertakings, there is an emerging trend for bottom-up initiatives, starting with small-scale projects built upon existing collaborations and infrastructures. As described by Mainen and colleagues (Mainen et al., 2016), these initiatives are centralized around self-organized groups of researchers working on the same challenges and sharing interests and specialized expertise. These projects could scale and open up to a larger audience and other disciplines over time, eventually lining up and merging their findings with other programs to make the bigger picture.

### 10Kin1day

One type of event that fits well with this grass-roots collaboration philosophy are short gatherings of scientists around a single theme, bringing together expertise and tools to jointly analyze existing neuroscience data. We organized 10Kin1day, an MRI connectome event, with the goal to bring together an international group of researchers in the field of neuroimaging and consistently analyze MRI connectivity data of the human cerebrum. We organized the event around five founding principles:

- use existing neuroimaging data, available from many research groups around the world; we focused on diffusion MRI data and aimed to bring together 10,000+ datasets
- analyze data from varying cohorts and imaging protocols, using a single, straightforward analysis strategy to encourage across-group collaborations and multisite studies
- perform all processing during a short workshop, with only basic expertise of analysis needed
- provide education on how to analyze resulting connectome data, so participants can continue to work on their projects after the event
- each participant analyzes their own data and is free to decide what to do with their analyzed results

### The 10K workshop

Over 50 participants from 40 different neuroimaging groups gathered in The Netherlands for a 3-day event. Participants brought and worked on their own datasets, varying from MRI data on healthy human brain organization, cross-sectional and longitudinal brain development, aging, cognitive psychology, as well as MRI data of a wide range of neurological and psychiatric brain disorders (including among others: Schizophrenia, Mood Disorders, Alzheimer’s Disease, Mild Cognitive Impairment, Amyotrophic Lateral Sclerosis, Frontotemporal Dementia, Epilepsy and Parkinson’s Disease). Written informend consent of the included healthy controls and/or patients was obtained by each of the participants at their local institute. 10 TB online storage space and 50,000+ CPU hours was reserved on the Cartesius supercomputer of the collaborative Information and Communication Technology (ICT) organization for Dutch education and research (SURF, https://surfsara.nl/) to analyze the data during the workshop. Workshop participants performed data quality checks on their data one week before the event after which they uploaded the MRI data (Diffusion Weighted Images (DWI) and pre-processed T1 data, see Materials and Methods) to their own user account on the supercomputer. During the workshop, participants were brought up to speed on DWI processing, connectome construction (see methods for details on the performed analysis), and running parallel jobs on a supercomputer. Together, a total of 15,947 MRI datasets were processed into anatomical connectome maps, with each output dataset including connectivity matrices with different types of connection weights and multiple parcellation resolutions. Data processing was paralleled by interactive educational talks and workshops on connectome analysis.

### Open data

In line with the collaborative nature of the event, the 10K group discussed making the connectome maps available to the scientific community for non-commercial use, free of restrictions. We include herein the resulting individual connectome maps of 8,000+ connectome datasets across an age range of 0 - 90 years, with five different edge weights (number of traced streamlines (NOS), streamline density (SD), fiber length, fractional anisotropy (FA) and mean diffusivity (MD)) at three parcellation resolutions (80+ cortical and subcortical regions, 100+ and 200+ cortical regions, see methods for details). Connectome maps are presented anonymously and blinded for participation site, together with basic demographics (age in bins of 5 years, gender, patient/control status, Fig. 1). Data is presented under the Non-Commercial Common Creative (CC BY-NC) license, free for all scientists to use in a non-commercial setting. A download request can be made at dutchconnectomelab.nl/10Kdata for a download link to the data (including 3 atlas resolutions, connectome matrices with multiple weights, information on the cortical and subcortical nodes, subject demographics (gender, age in 5 year bins, case/control)).

**Figure 1.**
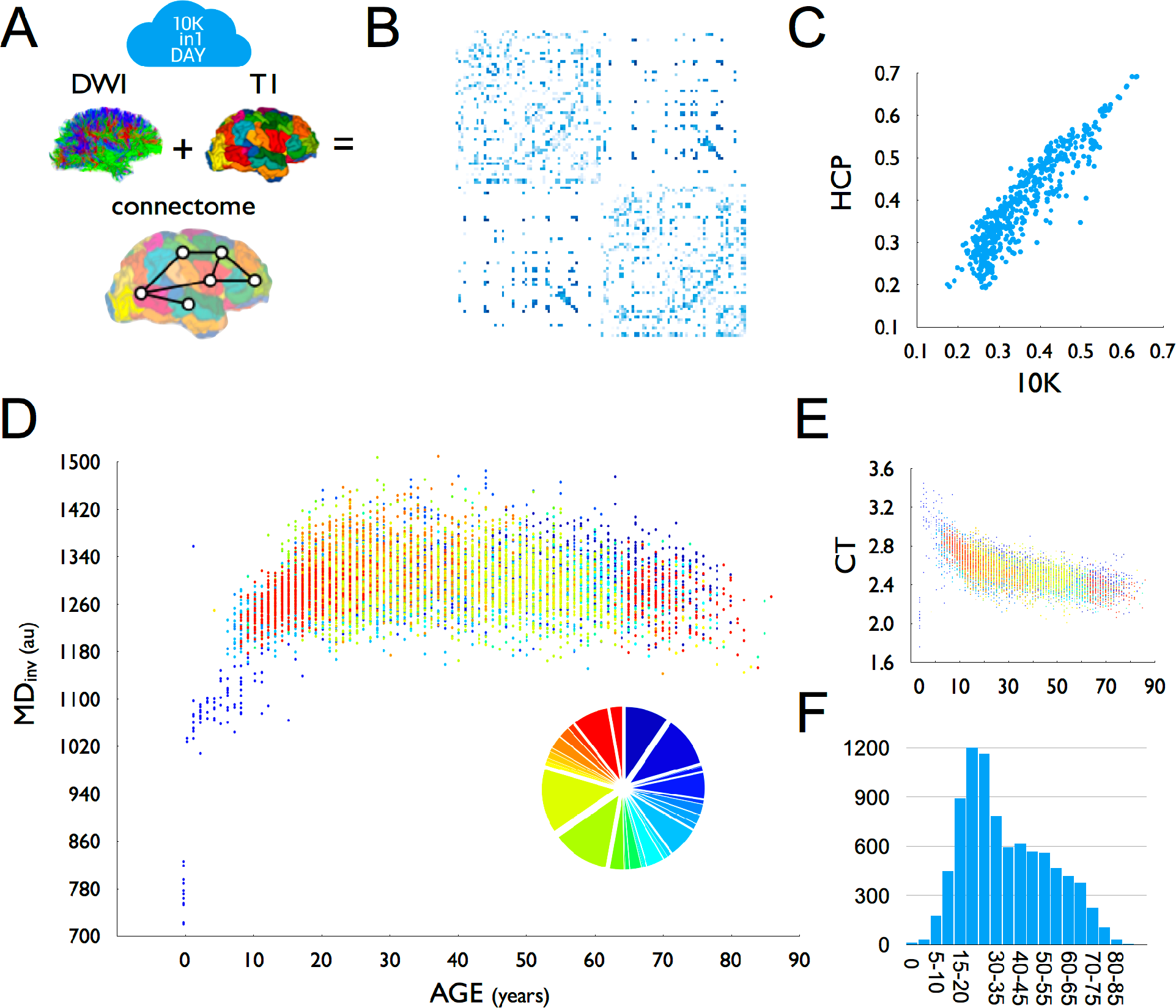
(A) For each dataset, DWI tractography was combined with T1-based parcellation of cerebral brain regions to reconstruct a brain network. (B) Group-averaged (group threshold 33%) FA matrix of the 10K dataset. (C) High overlap (r=0.93) between group-averaged FA values as derived from high-resolution HCP data and the 10K dataset. (D) Relationship between age and average inverse mean diffusivity (MD) across the 10K dataset. Colors indicate the different included datasets. Insert shows a pie diagram of the size of included datasets, color coded to set participation. One dataset (set_634413) was excluded from this plot, showing (across the age span) deviating FA (lower) and MD (higher) values than the other datasets (see methods). Due to the high total n, excluding this dataset did not change the relationship with age. (E) Relationship between age and average cortical thickness (CT). (F) Age distribution of the presented data as in panel E and F. au=arbritrary units. DWI=diffusion weighted imaging. CT=cortical thickness.

### A first peek

We performed a few first analyses on the joint dataset, such as comparison to Human Connectome Project (HCP) data and examination of effects of age (see Materials and Methods for more detail). We observed a high consistency of the group averaged matrix with data derived from the high-quality HCP, with at least 69% of pathways identified in HCP also observed in the 10K set and with 98% of all non-existing connections in HCP verified in the 10K set. Furthermore, the distribution of weights across reconstructed connections is highly similar across the two datasets (FA weights, r=.93, p<0.0001, Fig. 1). Inverse MD showed rapid growth of microstructure in early years, with continuing development throughout adolescence, peaking around the beginning of the third decade, followed by a steady pattern of decline throughout aging (Fig. 1). Age analysis of the 10K set shows clear developmental patterns of cortical morphology (Fig. 1 showing cortical thickness) and white matter microstructure (Fig. 1 showing inverse MD) across age.

### Future events

We acknowledge that there are many shortcomings to the presented MRI connectome dataset. Besides general, inherent limitations of diffusion MRI (Mori and Zhang, 2006), the presented dataset is a collation of data from a wide variety of groups, acquired with different scanners, different scanning protocols, varying data quality et cetera, as well as including data from a mixture of different patient and control populations. While these limitations place constraints on the type of investigations that one can perform with such collated multi-site datasets, we are optimistic that the 10K dataset can be used as a large reference dataset for future studies, enabling many technical and neuroscientific research questions to be addressed (e.g. Fig. 1). As such, we hope that the presented data will be of use to the neuroscience community in the examination of the human connectome. Above all, we hope that our report will inspire others to organize exciting 10Kin1day-type of events in the near future, bringing together existing neuroimaging data and further catalyze open neuroimaging research of the healthy and diseased brain.

## Author contributions

All authors contributed to the acquisition and/or analysis of the MRI data. Author Martijn van den Heuvel wrote the manuscript. All authors revised the manuscript and contributed intellectual content.

## Acknowledgements

The 10Kin1day workshop was generously sponsored by the Neuroscience and Cognition program Utrecht (NCU) of the Utrecht University (https://www.uu.nl/en/research/neuroscience-and-cognition-utrecht), the ENIGMA consortium (http://enigma.ini.usc.edu), and personal grants: Martijn P. van den Heuvel: NWO-VIDI (452-16-015), MQ 2015 Fellowship; Simon Baron-Cohen: the Wellcome Trust; Medical Research Council UK; NIHR CLAHRC for Cambridgeshire and Peterborough Foundation National Health Services Trust; Autism Research Trust; Linda Booij: New Investigator Award, Canadian Institutes of Health Research; Dara Cannon: Health Research Board (HRB), Ireland (grant code HRA-POR-2013-324); Sandra Chan: Research Grant Council (Hong Kong)-GRF 14101714; Eveline Crone: ERC-2010-StG-263234; Udo Dannlowski: DFG, grant FOR2107 DA1151/5-1, SFB-TRR58, Project C09, IZKF, grant Dan3/012/17; Stefan Du Plessis: MRC-RFA-UFSP-01-2013 (“Shared Roots” MRC Flagship grant); Thomas Frodl: Marie Curie Programme, International Training Programme, r’Birth; Dariusz Gąsecki: National Science Centre (UMO-2011/02/A/NZ5/00329); Beata Graff: National Science Centre (UMO-2011/02/A/NZ5/00329); Julie Hall: Western Sydney University Postgraduate Research Award; Laurena Holleran: Science Foundation Ireland, ERC; Helene Hopman: Research Grant Council (Hong Kong)-GRF 14101714; Lutz Jäncke: Velux Stiftung, grant 369; Andreas Jansen: DFG, grant FOR2107 JA 1890/7-1; Krzysztof Jodzio: National Science Centre (UMO-2013/09/N/HS6/02634); Vasiliy Kaleda: The Russian Foundation for Basic Research (grant code 15-06-05758 A); Tilo Kircher: DFG, grant FOR2107 KI 588/14-1, DFG, grant FOR2107 KI 588/15-1; Axel Krug: DFG, grant FOR2107 KO 4291/4-1, DFG, grant FOR2107 KO 4291/3-1; Irina Lebedeva: The Russian Foundation for Basic Research (grant code 15-06-05758 A); Edwin Lee: Health and Medical Research Fund - 11121271; Simon Lewis: NHMRC-ARC Dementia Fellowship 1110414, NHMRC Dementia Research Team Grant 1095127, NHMRC Project Grant 1062319; Carlos Lopez-Jaramillo: 537-2011, 2014-849; Andrew McIntosh: Wellcome Trust Strategic Award (104036/Z/14/Z); Christian Montag: Heisenberg-Grant, German Research Foundation, DFG MO 2363/3-1; Pedro Moreira: Foundation for Science and Technology, Portugal - PDE/BDE/113601/2015; Krzysztof Narkiewicz: National Science Centre (UMO-2011/02/A/NZ5/00329); Patrycja Naumczyk: National Science Centre (UMO-2013/09/N/HS6/02634); Jiska Peper: NWO-Veni 451-10-007; Paul Rasser: PER and US would like to thank the Schizophrenia Research Institute and the Chief-Investigators of the Australian Schizophrenia Research Bank V. Carr, U. Schall, R. Scott, A. Jablensky, B. Mowry, P. Michie, S. Catts, F. Henskens, and C. Pantelis.; Agnieszka Sabisz: National Science Centre (UMO-2011/02/A/NZ5/00329); Stavros Skouras: European Union’s Horizon 2020 research and innovation programme under the Marie Skłodowska-Curie grant agreement No 707730; Carles Soriano-Mas: Carlos III Health Institute (PI13/01958), Carlos III Health Institute (PI16/00889), Carlos III Health Institute (CPII16/00048); Edyta Szurowska: National Science Centre (UMO-2011/02/A/NZ5/00329); Alexander Tomyshev: The Russian Foundation for Basic Research (grant code 15-06-05758 A); Diana Tordesillas-Gutierrez: PI14/00918, PI14/00639; Leonardo Tozzi: Marie Curie Programme, International Training Programme, r’Birth; Sofie Valk: IMPRS Neurocom stipend; Theo van Erp: National Center for Research Resources at the National Institutes of Health (grant numbers: NIH 1 U24 RR021992 (Function Biomedical Informatics Research Network), NIH 1 U24 RR025736-01 (Biomedical Informatics Research Network Coordinating Center; http://www.birncommunity.org) and the NIH Big Data to Knowledge (BD2K) award (U54 EB020403 to Paul Thompson). Neeltje van Haren: NWO-VIDI (452-11-014); Judith van Leeuwen: NWO Veni (451-13-001); Marta Witkowska: National Science Centre (UMO-2011/02/A/NZ5/00329); Veronica O’Keane: Meath Foundation. The funding sources had no role in the study design, data collection, analysis, and interpretation of the data.

HCP data was provided by the Human Connectome Project, WU-Minn Consortium (Principal Investigators: David Van Essen and Kamil Ugurbil; 1U54MH091657) funded by the 16 NIH Institutes and Centers that support the NIH Blueprint for Neuroscience Research; and by the McDonnell Center for Systems Neuroscience at Washington University.

## Supplementary Files

- Supplementary Table 1 describing the dataset demographics
- Group files (3 atlas resolutions) containing 8,000+ connectome maps. A link to download the connectome matrices (2 Gb) can be obtained at http://dutchconnectomelab.nl/10Kdata.

## Materials and Methods

A total of 42 groups (52 participants) participated in the workshop, some working on multiple datasets. Each dataset included a diffusion MRI scan and T1 MRI scan processed using FreeSurfer (Fischl and Dale, 2000). Datasets across groups included data from 1.5 and 3 Tesla MRI with varying scanner protocols and number of applied DWI gradients. Data included MRI data of healthy participants and patients with a neurological or psychiatric disorder. 24 groups were able to make their data available, making a total of 8,697 connectome maps publicly available through means of this report. Reconstructed connectome maps are presented anonymously, blinded for participation site and disease condition. Basic demographics of the datasets are included in the download set.

### DWI Preprocessing

DWI datasets were corrected for susceptibility and eddy current distortions using the open tools from the FMRIB Software Library (FSL, http://fsl.fmrib.ox.ac.uk). Depending on the included DWI dataset, participants could choose to preprocess their data using the FSL *eddy_correct* or *eddy* tool (preprocessing scripts are included as supplementary information). For those DWI sets that included a subset of scans with an opposite k-space read out, an additional field distortion map could be formed and applied to the DWI images (Andersson et al., 2003).

### Cortical parcellation

Before the event, the participants created FreeSurfer files based on their T1 images, with this output being subjected to varying degrees of quality control. The resulting parcellations of the cerebrum were used to select the regions of interest for the connectome reconstruction. The 68 cortical regions of FreeSurfer’s standard Desikan-Killiany Atlas (Desikan et al., 2006, Fischl et al., 2004) as well as 14 subcortical regions were selected as network regions. Additionally, FreeSurfer files were used to further parcellate the cortex into 114 and 219 regions respectively using the Cammoun atlas (Cammoun et al., 2012).

### Fiber reconstruction

After preprocessing of the DWI data, a diffusion tensor was fitted to the diffusion signal in each voxel of the white matter mask (selected based on the white matter segmentation map of the FreeSurfer files) using robust tensor fitting (Chang et al., 2005). Simple Diffusion Tensor Imaging (DTI) reconstruction was used due to its robustness and relatively low sensitivity to false positive reconstructions compared to more advanced reconstruction methods (Klaus Maier-Hein, 2017), and thus potentially being the least distorting solution for connectome reconstruction and analysis based on MR imaging data (Zalesky et al., 2016). Decomposition of the tensor into eigenvectors and eigenvalues was used to select the main diffusion direction in each voxel, and to compute fractional anisotropy (FA) and mean diffusivity (MD) (Beaulieu and Allen, 1994). Deterministic fiber tractography was used to construct large-scale white matter pathways. Eight seeds (evenly distributed across the voxel) started in each white matter voxel, and fiber streamlines were formed by following the main diffusion direction from voxel to voxel using the *fiber assignment by continuous tracking* (FACT) algorithm (Mori and Barker, 1999), until one of the stopping criteria was met. A streamline was stopped when (1) it hit a voxel with an FA<0.1, (2) went out of the brain mask, or (3) made a turn >45 degrees.

### Connectome reconstruction

A connectome map was made by combining the (sub)cortical parcellation map and the set of reconstructed fibers using commonly described procedures (Hagmann et al., 2008; van den Heuvel et al., 2012; van den Heuvel et al., 2010; van den Heuvel and Sporns, 2011). For each of the Cammoun Desikan-Killiany parcellation maps (i.e. 14+68, 14+114 and 14+219 regions respectively), the total collection of reconstructed fiber streamlines was used to assess the level of connectivity between each pair of (sub)cortical regions, represented as the *connectivity matrix CIJ*. (Sub)cortical regions were selected as the nodes of the reconstructed network, and for each combination of region *i* and region *j* where fiber streamlines touched both regions a connection (i.e. network edge) was included in cell *CIJ(i,j)* in the connectivity matrix. Five different types of strength of a connection were computed and included as edge strength: (1) the number of reconstructed streamlines (NOS) between region *i* and *j*, (2) the average FA of the voxels traversed by the reconstructed streamlines, (3) the average MD of the reconstructed streamlines, (4) the average length of the reconstructed streamlines and (5) streamline density computed as the number of reconstructed streamlines corrected for the average volume of region *i* and region *j* (Hagmann et al., 2008).

### Outliers

A total of 15,947 connectome maps were analyzed across the participating groups. Of the datasets that could be shared, 197 were detected as outliers (and were subsequently removed from the dataset). Outliers were detected automatically per group by testing for each connectome map their average connection strength and their distance to the group average prevalence map. The average connection strength of a connectome map was calculated for each of the five connection weights as the mean of the strengths over all existing (nonzero) connections. To measure the presence of odd connections or absence of common connections in a connectome map, we constructed a group prevalence matrix for each dataset, counting per node pair how many times an edge was observed across connectome maps in the group. For each connectome map the total prevalence of all observed connections and the total prevalence of all non observed connections was computed. Outliers were identified as connectome maps that displayed on any of the 7 measures (5 weight and 2 prevalence measures) a score below Q1 – 2×IQR or above Q3 + 2×IQR, with Q1 and Q3 referring to the first and third quartile respectively and IQR the interquartile range IQR = Q3 – Q1. This resulted in the detection of 197 outliers in total, which were excluded from the dataset. One complete dataset (set_634413, n =584) showed across all included individual sets an average lower FA / higher MD as compared to the other datasets and this set was excluded from the age curves shown in Figure 1. Due to the high overall sample size, including or excluding this dataset did not change the shape of the final plot.

### Comparison to HCP data

To test the validity of the 10K dataset, we compared the group average matrix of the 10K set to the group average matrix of data from the Human Connectome Project (HCP) (Van Essen et al., 2013). First, for the 10K dataset, a group average FA matrix was computed, by including those edges that were observed in at least 33% of the group (i.e. a group threshold of 33%, >2700 connectome maps showing a particular network edge). Average weight values of the included edges were taken as the non-zero mean of those edges across the connectome maps. Second, a similar group average FA matrix was derived from previously analyzed HCP data (van den Heuvel et al., 2016) (n=487 datasets). In brief, HCP analysis included the following steps (see (van den Heuvel et al., 2016) for more detailed information on the HCP data analysis). For each of the HCP DWI datasets a connectome was reconstructed based on the minimally pre-processed data of HCP. Given the high quality of the HCP data, analysis here included reconstruction of multiple diffusion directions, allowing for the reconstruction of more complex fiber configurations (e.g. crossing fibers) (van den Heuvel et al., 2016). Similarly as for the 10K data, across the total set of 487 datasets, an average FA group matrix was computed, including those network edges that were observed in at least 33% of the total population (i.e. >160 datasets) and taking the non-zero mean of FA values across the group of subjects. Comparison between the 10K set and the HCP dataset was computed by means of (1) counting the number of existing connections and non-existing connections in the 10K dataset as observed in the HCP dataset and (2) by correlating the FA weights of the set of edges as observed in both datasets.

